# Habitat generalists or specialists, insights from comparative genomic analyses of *Thermosipho* lineages

**DOI:** 10.1101/106989

**Authors:** Thomas H.A. Haverkamp, Claire Geslin, Julien Lossouarn, Olga A. Podosokorskaya, Ilya Kublanov, Camilla L. Nesbø

## Abstract

*Thermosipho* species inhabit various extreme environments such as marine hydrothermal vents, petroleum reservoirs and terrestrial hot springs. A 16S rRNA phylogeny of available *Thermosipho* spp. sequences suggested habitat specialists adapted to living in hydrothermal vents only, and habitat generalists inhabiting oil reservoirs, hydrothermal vents and hotsprings. Comparative genomics and recombination analysis of the genomes of 15 *Thermosipho* isolates separated them into three species with different habitat distributions, the widely distributed *T. africanus* and the more specialized, *T. melanesiensis* and *T. affectus*. The three *Thermosipho* species can also be differentiated on the basis of genome content. For instance the *T. africanus* genomes had the largest repertoire of carbohydrate metabolism, which could explain why these isolates were obtained from ecologically more divergent habitats. The three species also show different capacities for defense against foreign DNA. *T. melanesiensis* and *T. africanus* both had a complete RM system, while this was missing in *T. affectus*. These observations also correlated with Pacbio sequencing, which revealed a methylated *T. melanesiensis* BI431 genome, while no methylation was detected among two *T. affectus* isolates. All the genomes carry CRISPR arrays accompanied by more or less complete CRISPR-cas systems. Interestingly, some isolates of both *T. melanesiensis* and *T. africanus* carry integrated prophage elements, with spacers matching these in their CRISPR arrays. Taken together, the comparative genomic analyses of *Thermosipho* spp. revealed genetic variation allowing habitat differentiation within the genus as well as differentiation with respect to invading mobile DNA that is present in subsurface ecosystems.

## Introduction

Bacteria of the genus *Thermosipho* belong to phylum Thermotogae and are found in high temperature environments such as deep-sea hydrothermal vents, and subsurface oil reservoirs (Antoine et al. 1997; Takai & Horikoshi 2000; Dahle et al. 2008). There are currently eight described *Thermosipho* species: *T. africanus*, *T. activus*, *T. affectus*, *T. atlanticus*, *T. geolei*, *T. globiformans*, *T. japonicus* and *T. melanesiensis* (Mukherjee & Sengupta 2015). These are thermophilic, organotrophic, anaerobes fermenting various sugars of different complexity (e.g. glucose, starch, cellulose, etc) and peptides. While the main fermentation products of all species are H_2_ and CO_2_, species-specific production of compounds such as acetate, lactate, ethanol or alanine have been described (Swithers et al. 2011).

Comparative genomic analyses of Thermotogae bacteria, including *Thermosipho*, have revealed complex evolutionary histories with extensive horizontal gene transfer (HGT), particularly involving members of the bacterial phylum Firmicutes and the Archaea domain (Enright et al. 2002). Most of the unique genes with predicted functions in different Thermotogae lineages are classified as being involved in metabolism and particularly genes involved in carbohydrate transport and degradation are numerous (Nesbø et al. 2002; Zhaxybayeva et al. 2009). In agreement with this, Thermotogae and *Thermosipho* species in particular, are commonly distinguished by being able to grow on different substrates. For instance, *T. affectus* and *T. activus* are able to degrade cellulose compounds, while others cannot (e.g. *T. africanus*) (Podosokorskaya et al. 2014). *T. japonicus* is able to degrade casein as the sole carbon source in conjunction with the electron acceptor thiosulfate (Na_2_S_2_O_3_) (Takai & Horikoshi 2000). How such metabolic differences are encoded in the genomes and what pathways are involved is unresolved. Furthermore, how do the genetic differences correlate with the diversity found in the *Thermosipho* genus and the ecosystems that they occupy?

One feature of the *Thermosipho* genus that has been studied at the genome level is the presence of the vitamine B12 synthesis pathway in several *Thermosipho* isolates, but is absent in other Thermotogae (Swithers et al. 2011). The genes needed for the vitamin B12 pathway were acquired horizontally from the phylum Firmicutes. The acquisition of novel genetic material in this way allows prokaryotes to establish novel metabolic capacities and develop specific adaptations which may lead to species differentiation and innovation (Koonin 2015). In addition, HGT is suggested to be important in the long-term maintenance and repair of genomic information, by replacing inactivated genes by active ones (Takeuchi et al. 2014). HGT can take place via transformation (naked DNA), conjugation (plasmids), transduction (viruses), or transposition (transposable elements) but also with gene transfer agents (GTA’s), nanotubes or membrane vesicles produced by the microorganisms themselves (Darmon & Leach 2014).

Although novel DNA may be beneficial to microorganisms (Darmon & Leach 2014), viruses (and many mobile elements) can be without doubt harmful for cells, and prokaryotes have developed defense methods. Several mechanisms exist that regulate the activity of mobile DNA inside the cell (Westra et al. 2012), such as the restriction-modification (RM) system, abortive infection (ABi) mechanisms, the clustered regulatory interspaced short palindromic repeats (CRISPR) system and the recently described bacteriophage exclusion (BREX) system (Stern & Sorek 2011; Goldfarb et al. 2015). CRISPRs seems to be particularly important in thermophiles (Weinberger et al. 2012), and have been identified in all Thermotogae genomes characterized to date including two *Thermosipho* genomes (Nesbø et al. 2009; Zhaxybayeva et al. 2009). The CRISPR associated genes (cas-genes) are essential for the adaptation step in the CRISPR defense system where spacers from invading DNA are acquired and inserted in the CRISPR array (Makarova et al. 2015). Interestingly, most of the CRISPR associated genes (Cas) show evidence of HGT when comparing *T. africanus* to *Thermotoga maritima* (Nesbø et al. 2009).

The well-studied bacterial RM system is almost universally found in both bacteria and archaea (Roberts et al. 2010). The RM system uses methylation to distinguish between self and foreign DNA, and is found in most bacterial genomes (Roberts et al. 2010; Vasu & Nagaraja 2013). The system works by protecting methylated DNA from restriction enzymes. Those enzymes will therefore only degrade invading mobile DNA that is not methylated. Although the RM system is well studied, it is only poorly characterized among the Thermotogae species described so far (Xu et al. 2011). For the genus *Thermosipho* in particular, it is still unclear if RM systems are present, functional, and how it is involved in the maintenance of genome stability when these populations have a high quantity of HGT acquired genes.

In light of the high numbers of horizontally acquired genes within the Thermotogae it is interesting that within the species *Thermotoga maritima* we found highly similar genomes (Nesbø et al. 2015). This is remarkable when considering the isolation sources of the different strains (The Azores, Italy, Japan, Kuril Islands (Russia) and the North Sea), which span the earth, and suggests high levels of gene flow between distant subpopulations and active mechanisms to maintain highly similar genomes (Nesbø et al. 2015).

Here we present a comparative analysis of 15 *Thermosipho* genomes, thirteen of which are novel sequences generated in this study. The isolates included in this analysis were obtained from deep-sea hydrothermal vents and produced fluids from oil reservoirs. The genomes fall into three well-defined lineages (or species) with isolates from the same sample sites showing very high similarities to each other. We show that the three species differ in genomic content with regard to metabolic systems involved in carbohydrate and coenzyme metabolism, CRISPR-cas and the RM system. In addition, we show that several of the genomes contain prophages.

## Materials & Methods

### DNA extraction and genome sequencing

13 *Thermosipho* strains were cultured in a modified Ravot medium as previously described in (Lossouarn et al. 2015) and used for total DNA extraction as described in (Charbonnier et al. 1992; Geslin et al. 2003). All strains, except *T. africanus* Ob7 were submitted for genome sequencing at the Norwegian Sequencing Centre (NSC, Oslo, Norway). A 300 bp insert paired end library was prepared using the genomic DNA sample prep kit (Illumina, San Diego, CA, USA) for each DNA sample. The Library was spiked with phiX DNA. Subsequently, all strains except Ob7 were sequenced on an Illumina MiSeq generating a dataset of 250 bp paired end reads.

Strain Ob7 was sequenced at University of Alberta, Canada, using an Iontorrent (Thermo Fisher Scientific, Waltham, MA, USA) approach. DNA was enzymatically sheared using the Ion Shear Plus kit (Life Technologies, Carlsbad, CA, USA) then cloned using the Ion Plus Fragment Library kit (Life Technologies) following the manufacturer’s instructions. The library was then sequenced on an Ion Torrent PGM using a 316D chip and 500 flows.

Four isolates (*Thermosipho* sp. 430, 431, 1063, 1070) were selected for long read sequencing with the goal to complete the genomes and assess methylation status of the DNA. DNA’s were prepared following the Pacific Biosciences instructions and sequenced at the NSC on a Pacbio RSII (Pacific Biosciences, Menlo Park, CA, USA).

### Quality control and genome assembly

Pacbio sequencing of the strains 431, 1063 and 1070 produced libraries suitable for genome assembly without the addition of Illumina paired-end reads. For all strains Pacbio read filtering and error-correction were performed using version 2.1 of the Pacbio smrtportal (http://www.pacb.com/devnet/). Assembly and base modification detection were performed using the RS_HGAP_Assembly.2 and RS_Modification_and_Motif_analysis protocols.

Overlapping regions between contigs were manually detected using a combination of read mapping and subsequent visualization with IGV (v. 2.3.23) (Thorvaldsdottir et al. 2013). This allowed us to inspect regions with low quality mappings (due to the presence of duplicated regions), and similar gene annotations on both sides of a contig gap. Preliminary annotations were prepared using Glimmer at the RAST server and inspected in the CLC Bio main workbench (Overbeek et al. 2013). Bam files were generated by mapping MiSeq PE reads using bwa version (v. 0.7.5a) to the annotated contigs (Li & Durbin 2009) and by converting sam files with Samtools (v. 0.1.19). Bam files were imported into the IGV viewer together with the annotated contigs (Li et al. 2009). This approach resulted in closed chromosomal sequences.

For strain 430, we generated a combined assembly using high quality illumina MiSeq reads and a smaller set of Pacbio reads (15594 subreads) in an assembly using Spades (v. 3.5.0.) (Bankevich et al. 2012) with kmer size set to 127. Low coverage contigs (< 2.0) were discarded. The remaining contigs were checked in IGV for miss-assemblies using mapped MiSeq and Pacbio reads and if detected contigs were discarded. Finally the contigs were checked for unambiguously overlapping ends as described above and if detected the contigs were combined.

The *T. africanus* Ob7 genome was assembled using NEWBLER v. 2.6 (Margulies et al. 2005). The remaining strains, were assembled using CLC assembler v.7, Spades (v. 3.5.0.) and Velvet (v 1.2.10) (Zerbino & Birney 2008) (Table 1). Genome assemblies were compared using REAPR (v1.0.16) and the most optimal assembly for each strain was selected for genome annotation with the PGAP pipeline from NCBI (Hunt et al. 2013).

**Table 1:** Overview of *Thermosipho* genomes used in this study.

| T. genome | NCBI accession number | Nr of contigs # | Genome size (bp) | GC content (%) | Sequence technology | Genome coverage § | Isolation source | References |
| --- | --- | --- | --- | --- | --- | --- | --- | --- |
| *T. melanesiensis* BI429*& | NC_009616 | 1 | 1.915.238 | 31.4 | 454 / Sanger | - | Gills of mussel, Deep-sea H. vent, Pacific Ocean, Lau Basin | Antoine et al., 1997<br>Zhaxybayeva et al., 2009 |
| *T. melanesiensis* 430& | JYCX000000000 | 4 | 1.906.053 | 31.5 | MiSeq | 243<sup>M</sup> | Gills of gastropod Deep-sea H. |  |
| *T. melanesiensis* 431& | CP007389 | 1 | 1.915.344 | 31.4 | Pacbio /MiSeq | 34<sup>P</sup>/223<sup>M</sup> | Sample of active chimney | this study |
| *T. melanesiensis* 432& | JYCY000000000 | 21 | 1.890.294 | 32.6 | MiSeq | 230<sup>M</sup> | Sample of active chimney | this study |
| *T. melanesiensis* 433& | JYCZ000000000 | 21 | 1.890.575 | 32.6 | MiSeq | 256<sup>M</sup> | Gills of a mussel |  |
| *T. melanesiensis* 434& | JYDA000000000 | 22 | 1.892.506 | 32.7 | MiSeq | 234<sup>M</sup> |  |  |
| *T. melanesiensis* 487& | JYDB000000000 | 21 | 1.890.479 | 32.6 | MiSeq | 239<sup>M</sup> | Sample of active chimney |  |
| *T. affectus* 1063<sup>%</sup> | CP007223 | 1 | 1.766.633 | 31.5 | Pacbio /MiSeq | 38<sup>P</sup>/304<sup>M</sup> | Deep sea H. vent, Atlantic ocean | Wery et al., 2002 |
| *T. affectus* 1070<sup>%</sup> | CP007121 | 1 |  | 31.5 | Pacbio /MiSeq | 37<sup>P</sup>/257<sup>M</sup> | Menez-Gwen hydrothermal field | this study |
| *T. affectus* 1074<sup>%</sup> | LBEX000000000 | 20 | 1.788.307 | 33.3 | MiSeq | 258<sup>M</sup> | Menez-Gwen hydrothermal field | this study |
| *T. affectus* ik275mar | LBFC000000000 | 27 | 1.771.018 | 32.6 | MiSeq | 256<sup>M</sup> | Rainbow H.vent field | Podosokorskaya et al., 2011 |
| *T. affectus* sp. 1223<sup>°</sup> | LB EY000000000 | 39 | 1.767.555 | 32.1 | MiSeq | 226<sup>M</sup> | Rainbow H.vent field | this study |
| *T. africanus* ob7 | ? | 23 | 1.933435 | 32.7 | Ion Torrent | - | Tadjoura gulf Hydrothermal springs | Huber et al., 1989 |
| *T. africanus* H17ap60334 | AJIP0100000 | 49 | 2.083.551 | 29.9 | 454 / Sanger | 310 | production water Hibernia oil platform | this study |
| *T. africanus* TCF52B* | NC_011653 | 1 | 2.016.657 | 30.8 | Sanger | - | production water Troll C oil platform, North Sea | Dahle et al., 2008<br>Nesbø et al., 2009 |

\* Not sequenced during this study
& Strains 429 to 487 isolated during Oceanographic cruise Biolau: Pacific Ocean, Lau Basin, Deep-sea Hydrothermal vent .
<sup>%</sup> Strains 1063 to 1074 isolated during Oceanographic cruise Marvel: Atlantic ocean, Menez Gwen, Deep-sea Hydrothermal vent
<sup>∞</sup> Isolated during Oceanographic cruise Atos, Atlantic ocean, Rainbow, Deep sea Hydrothermal vent.
<sup>#</sup> Genomes with 1 contig were closed
<sup>§</sup> <sup>M</sup>: MiSeq sequencing; <sup>P</sup>: Pacbio RSII sequencing;

### 16S rRNA Phylogeny of Thermosipho species

Full-length 16S rRNA sequences were manually extracted from the annotated *Thermosipho* genome assemblies generated in this study as well as publically available genomes. Next we searched the SILVA database version SSU r122 for *Thermosipho* spp. 16S rRNA sequences with the following setting: sequence length > 1300 bp, alignment quality > 80, pintail quality > 80 (Quast et al. 2013). Matching sequences were downloaded and added to extracted 16S rRNA sequences from the de-novo sequenced genomes. Multiple identical sequences from the same strain with different accession numbers (e.g. the four 16S rRNA sequences of *T. melanesiensis* BI429, five 16S rRNA sequences *T. africanus* TCF52B) are represented by a single sequence in the dataset.

The NCBI non-redundant database was then queried using BLASTn (blast+ v 2.2.26 (Camacho et al. 2009)) with the 16S rRNA gene dataset described above, to identify *Thermosipho* sp. like 16S rRNA sequences missed with the above methods. Additional full-length sequences were downloaded from Genbank and combined with the de-novo / SILVA 16S rRNA sequences.

The final 16S rRNA gene alignment consisted of *Thermosipho* sequences combined with outgroup sequences from other Thermotogales species. The resulting alignment was used to build phylogenies using Maximum Likelihood as implemented in MEGA 6.0 (Tamura et al. 2013). and the General Time Reversible (GTR) model (G+I, 4 categories) as suggested by JModelTest v. 2.1.7 (Darriba et al. 2012).

### Pangenome analysis

Blast Ring genome plots were generated using BRIG version 0.95 (Alikhan et al. 2011) running Blast+ version 2.2.28 (Camacho et al. 2009) with the following set up. The nucleotide sequence of one of the 15 *Thermosipho* genomes was used to create the blast-database with default settings. Nucleotide sequences of all coding genes were extracted from each genome (NCBI Genbank annotation) using CLC Main workbench Version 6.8.3 and were used for BLASTn analysis in BRIG with the following settings: max_target_seqs: 1; max e-value cut-off: 1.0^−4^. Alignments with a minimum of 70% similarity were visualized with BRIG. The same procedure was used for all genomes.

Complete chromosome sequences or contigs were uploaded for all 15 genomes to the Panseq 2.0 server (https://lfz.corefacility.ca/panseq/) for pangenome analysis (Laing et al. 2010). Sequence similarity cut-off (SC) was set at 70 % to identify core-genome segments and SNP’s. We used the standard settings with Panseq except: percent sequence identity cut-off set to 70%, core genome threshold: 15, BLAST word size: 11. The final alignment of the single nucleotide polymorphisms (SNPs) was loaded into splitstree and visualized as an unrooted phylogeny using the neighbor network algorithm (Huson & Bryant 2006)

Ten genome sequences (none-closed genomes of the *T. melanesiensis* cluster were excluded, due to high sequence similarity) were aligned with progressiveMauve (Mauve (v. 2.3.1) (Darling *et al.*, 2010)) using *T. africanus* H17ap60344 set as the reference and automatically calculated seed weights and minimum Locally Colinear Blocks (LCB) scores. Gaps were removed and the edited LCBs were concatenated in Geneious 8 (www.geneious.com). Recombination analysis of the concatenated alignment was done in LikeWind (Archibald & Roger 2002a; 2002b) using the maximum likelihood tree calculated in PAUP* version 4.0b10 (Swofford 2002) under a GTR+Γ+Ι model as the reference tree.

Pairwise tetranucleotide frequency correlation coefficients (TETRA) and Average Nucleotide identity (ANI) were calculated using the JspeciesWS webtool (http://jspecies.ribohost.com/jspeciesws/) (Richter & Rosselló-Mora 2009; Goris et al. 2007). The pair-wise TETRA values were visualized with R (version 3.2.1) using the heatmap.2 function (Package gplots) and the *Thermosipho* genomes were clustered using Jaccard distances based on pairwise TETRA values (Package Vegan. version 2.3-0).

The IMG/ER Pairwise ANI calculation was used to determine the number of shared genes between each genome separately (Markowitz et al. 2009). The method uses pairwise bidirectional best nSimScan hits where similar genes share a minimum of 70% sequence identity with 70% coverage of the smaller gene. For each genome we calculated the fraction of shared genes with all other genomes. Unique genes per genome were determined using the IMG Phylogenetic profiler tool for single genes, where each genome was analyzed for the presence of genes without any homologs in all other 14 genomes. The settings were: minimum e-value of 1.0^-5^; min percent identity: 70%; pseudogenes were excluded; the algorithm was present/absent homologs. The same tool was used to identify the presence of homologs shared with all genomes, and with only the genomes from the same cluster.

### Functional comparison of genome

The 15 genomes were compared using the Clusters of Orthologous Genes (COGs) annotations. A COG reference database (version 10) was downloaded from the STRING database (http://string-db.org/). For each genome all protein sequences were aligned using BlastP, with the settings: 1 target sequence; maximum e-value: 1.0e^−20^; database size 1.0^7^; tabular output. Only hits to COG database sequences were retained when the alignment was >= 70% of the length of the longest protein sequence and were used to build a protein COG classification table. The COG IDs were summarized by classifying them to any of the available COG categories (ftp://ftp.ncbi.nih.gov/pub/wolf/COGs/COG0303/cogs.csv). For the COG analysis of the species-specific genes, we used the results from the IMG phylogenetic profiler tool to identify cluster specific genes. The COG classification table was screened for cluster specific genes and summarized into COG categories.

In order to statistically compare species differences of COG categories we normalized counts by total COG gene annotations, giving relative abundances per genome. R (version 3.3.0) was used to identify COG categories that were significantly different between the three species using the non-parametric Kruskal-Wallis test (p-value <= 0.01). The R-package ggplot2 (version 2.1.0) was used to generate the comparisons graphically.

### Horizontal gene transfer detection

Genes putatively acquired by HGT were identified using HGTector (Zhu et al. 2014) with BLASTp (blast+ v 2.2.26). The databaser.py script was used on 6 December 2015 to download per species one representative proteome of all microorganisms from the NCBI refseq database. We compared the predicted *Thermosipho* spp. protein sequences to the reference database using the following BLAST cut-offs: E-value: 1.0^−5^, Percentage identity: 70%, percentage coverage: 50%, and a maximum of 100 hits were returned. To determine which genes were putatively acquired by any of the strains we set the HGTector self group to the genus *Thermosipho* (NCBI taxonomy ID: 2420) and the close group to either the family Fervidobacteriaceae (NCBI taxonomy ID: 1643950), or the order Thermotogales (NCBI taxonomy ID: 2419). HGTector then analyzes the blast output from each protein for hits matching taxa belonging to either the self or close groups, or more distantly related taxa, which is used to determine which genes have likely been acquired from taxa more distantly related than the close group. When the close group was set to Fervidobacteriaceae, we identified putative HGT genes specific for *Thermosipo* sp. but not found in Fervidobacteriaceae species. This setting however, did not indicate putative HGT genes derived from other families within the Thermotogales order or beyond.

### Defense genes

Using the defense genes list created by Makarova et al., ( 2011), we screened the IMG genome Clusters of Orthologous Genes (COG) annotations for the presence of any COGs involved in any of the mobile DNA defences (Makarova et al. 2011). We summarized the identified COGs and distinguished between CRISPR-cas associated or restriction-modification (RM) system genes.

### Crispr-spacer analysis

All genomes were uploaded to CRISPR finder (http://crispr.i2bc.paris-saclay.fr/) to detect CRISPR-arrays (Grissa et al. 2007). CRISPR spacer sequences from each genome were compared using BLASTn against all *Thermosipho* sp. spacer sequences using the following settings: e-value cut-off: 1.0^−5^, database size: 1.0^7^, dust: no. The tabular blast results were visualized with R-statistics using the Markov Cluster Algorithm (MCL) (v1.0) and Igraph (v1.0.1) packages. Igraph was used for matrix construction. MCL was run using the matrix with the inflation set to 1.4 and max iterations set to 100 (Enright et al. 2002).

In order to search for matching sequences within the genome but outside the CRISPR arrays, e.g target genes, we masked all CRISPR arrays using maskfeat (EMBOSS v. 6.5.7 (Rice et al. 2000)). Next we ran BLASTn (v2.2.26+) using the spacers of each genome against the own genome using the following settings: e-value cut-off: 1.0^−5^, database size: 1.0^7^, dust: no. CRISPR array spacers were also compared against the NCBI nucleotide database to find other species with similar sequences. BLASTn was run with the settings: e-value cut-off: 1.0^−5^, dust: no. Each genome was screened for the presence of prophages (Supplementary information for details).

### Vitamine B_12_ pathway analysis

The genes involved in the Vitamine B_12_ metabolism are found in four different gene clusters (BtuFCD, Corriniod, Cobalamin, and SucCoA) in *Thermosipho* and can be regulated by B_12_ riboswitches (Swithers et al. 2011). All 15 genomes were screened for the presence of Cobalamin specific riboswitches using Riboswitch scanner (Mukherjee & Sengupta 2015). This information was used to confirm the presence of the four gene clusters in each genome. Next, we extracted the protein sequences from the *T. melanesiensis* BI429 genome involved in B_12_ metabolism (Swithers et al. 2011) and used them to identify homologous genes in all *Thermosipho* genomes using tBLASTn with a maximum e-value 1.0^-20^.

### Data desposition

All genomes were deposited in the Genbank database and their accession numbers are found in Table 1. In addition all genomes were deposited in the IMG databases and are linked to the NCBI accession numbers. The 16S rRNA alignment is available upon request from the corresponding author.

## Results

### Global isolation of the genus Thermosipho from hydrothermal vents and oilfields

We screened the NCBI-non-redundant (nr) database using BLASTn with *Thermosipho* 16S rRNA genes as probes to assess how our genomic analyses span the available environmental diversity of *Thermosipho* lineages. This analysis revealed that most lineages or species, e.g. *T. melanesiensis*, *T. affectus* and *T. africanus*, are well covered by our genomic analysis (Figure 1, Supplementary materials Figure S1). Nonetheless, there are several lineages, including *T. geolei*, *T. ferriphilus* and *T. activus*, for which we do not have genomic data yet.

**Figure 1.**
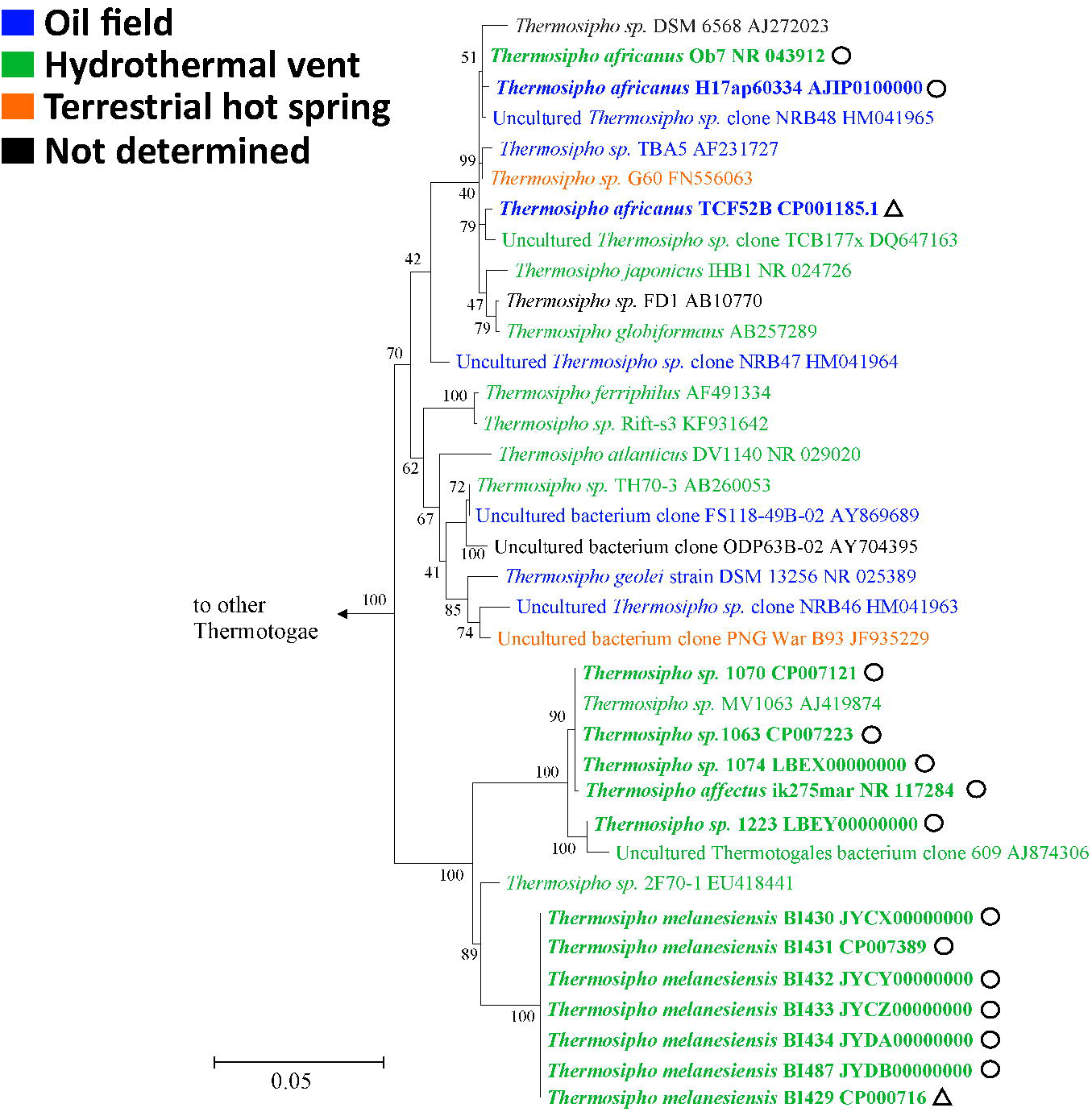
Maximum Likelihood phylogeny of *Thermosipho* 16S rRNA sequences constructed in conducted in MEGA6 (Tamura et al., 2013). The General Time Reversible (GTR) model (G+I, 4 categories, (Nei & Kumar 2000)) was used. The tree with the highest log likelihood is shown. The percentage of trees in which the associated taxa clustered together is shown next to the branches. Initial tree(s) for the heuristic search were obtained by applying the Neighbor-Joining method to a matrix of pairwise distances estimated using the Maximum Composite Likelihood (MCL) approach. The tree is drawn to scale, with branch lengths measured in the number of substitutions per site, bar indicates 0.05 substitutions per site. The analysis involved 61 nucleotide sequences. All positions with less than 95% site coverage were eliminated. That is, fewer than 5% alignment gaps, missing data, and ambiguous bases were allowed at any position. There were a total of 1245 positions in the final dataset. The phylogeny is rooted using 16S rRNA gene sequences of representative sequences from other Thermotogae genera (*Defluviitoga*, *Fervidobacterium*, *Geotoga*, *Kosmotoga*, *Marinitoga*, *Mesoaciditoga*, *Mesotoga*, *Oceanotoga*, *Petrotoga*, *Pseudothermotoga* and *Thermotoga*). All non-*Thermosipho* Thermotogales formed an outgroup and were collapsed (Supplementary figure S1 for the full phylogeny). *Thermosipho* sequences are colored based on the environment of isolation. Sequences with bold fonts are whole genome sequences. Triangles behind the sequence ID indicate genome not from this study. Circles behind sequence ID indicate genome from this study.

Moreover, the 16S rRNA gene phylogeny suggests ecological differences between the different lineages (Figure 1). *T. africanus* isolates appear to be habitat ‘generalists’ and have been isolated and detected in oil reservoirs, marine hydrothermal vents and terrestrial hot springs. In contrast the *T. melanesiensis* and *T. affectus* lineages appear to be more specialized and have only been obtained from marine hydrothermal vents. Thus one interesting question is if and how these different life styles are reflected in their genomes with regard to diversity and genome content.

### Genome overview

The current study added 13 new genomes, with different levels of completion, to the two existing ones (Table 1). All genomes share a low GC content (29.9 % − 32.7 %). Interestingly, the *T. affectus* genomes are the smallest in this dataset (≈ 1.77Mbp), while the T. *melanesiensis* (≈ 1.9Mbp) and *T. africanus* (≈ 2.0Mbp) genomes are larger.

### Phylogenomic analysis

Tetranucleotide frequencies can be used to calculate the genomic similarity between bacterial isolates, where pairwise similarity is expressed as tetranucleotide frequency correlation coefficients (TETRA) (Richter & Rosselló-Mora 2009). The heatmap of the TETRA values of the *Thermosipho* genomes indicated the presence of 3 groups with high intra relatedness (TETRA > 0.99) (Figure 2). Interestingly, the *T. africanus* isolates show more divergent genomes compared to the other two clusters. We obtained similar clustering results when we calculated the pairwise Average Nucleotide Identities (ANI) between the isolates (data not shown), with intracluster identities > 95% and intercluster identities < 90%. The core-and pangenome of the genus *Thermosipho* was estimated to be 1.4Mbp and 5.6Mbp, respectively (Panseq2 sequence identity cut-off (IC): 70%). Splitstree networks (Huson & Bryant 2006) using different SC showed very little differences regardless of which IC was used (data non shown). The unrooted neighbor network calculated from 427,560 core SNPs is shown in Figure 3A. The network shows three branches for the three lineages with few differences within each lineage, similar to the pattern shown in the 16S rRNA tree and the TETRA heatmap.

**Figure 2.**
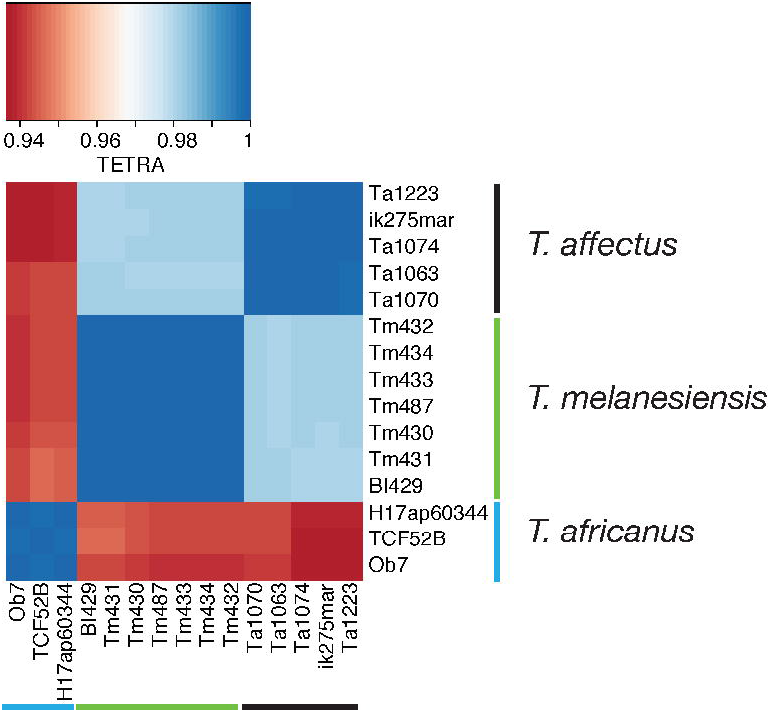
Comparison of the *Thermosipho* genomes based on pairwise tetranucleotide frequency correlation coefficients (TETRA). Dark blue indicate highly similar genomes, while red indicate low similarity. The colored lines (black, green and blue) indicate to which *Thermosipho* lineage/species the strains belong. The heatmap was created in R-studio using pairwise TETRA values calculated with the JSpeciesWS server (Richter et al., 2015).

Finally, we used the progressive Mauve aligner to extract Locally collinear blocks (LCBs) from 10 representative *Thermosipho* genomes to build a concatenated alignment. The phylogeny based on this alignment (Supplementary materials Figure S2A) showed a similar pattern as when using single nucleotide polymorphisms (SNPs) (Figure 3A). The concatenated Mauve alignment was used for recombination detection analysis using LikeWind (Archibald & Roger 2002b), which detected numerous recombination events within each cluster. However, no recombination events were detected between the clusters suggesting that the lineages do indeed correspond to three distinct species (Supplementary materials Figure S2).

**Figure 3.**
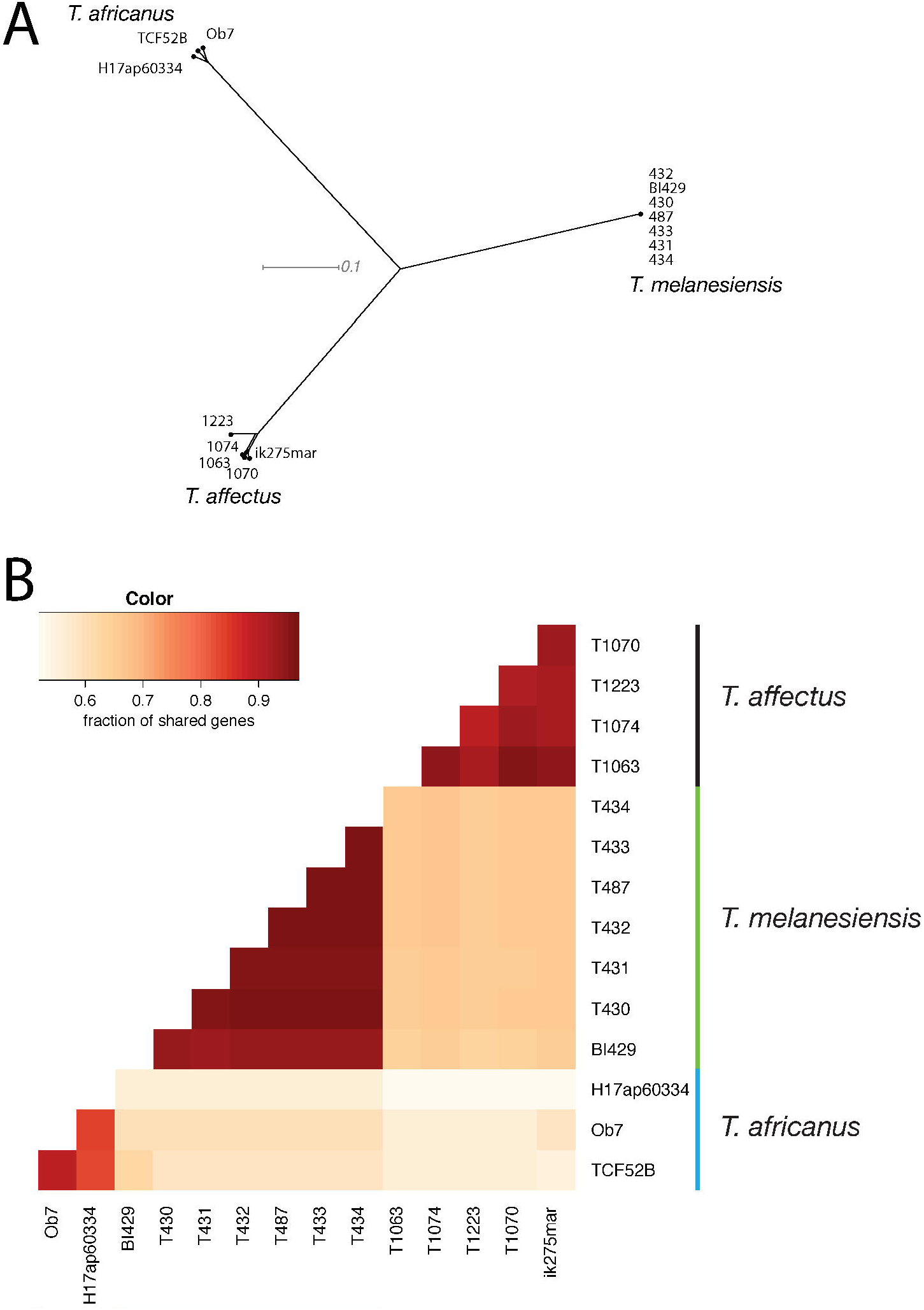
A) Neighbor network of the 15 *Thermosipho* strains. The network is based SNP’s in core genome fragments that were present in all genomes with a minimum of 70% similarity. The network was visualized in Splitstree using the NeighborNet algorithm (Huson and Bryant, 2006) from uncorrected distances. B) Fraction of genes shared between 15 *Thermosipho* genomes. Data was generated at the IMG-database using pairwise Bidirectional Best nSimScan Hits, with genes sharing 70% sequence identity and at least 70% coverage. Percentages were calculated by dividing the number shared genes by the total number of genes for the genome on the y-axis. The colored lines (black, green and blue) indicated which strains belong to which *Thermosipho* lineage/species.

### Core / Pan genes

Using the IMG Pairwise ANI tool we obtained for each genome the number of genes shared with all other genomes (Figure 3B). This gave a similar pattern as obtain using TETRA, ANIb/m, with the presence of three clusters. Between the three clusters we found that less than 80 % of the genes are shared between the species (IC: 70%). These results suggest the presence of strain and species-specific genes in each of the *Thermosipho* isolates analyzed. A visual inspection of genome content using BRIG showed a similar pattern (Supplementary information; Supplementary materials Figure S3 (Alikhan et al. 2011). The genomes from the *T. melanesiensis* cluster are highly similar, with *T. melanesiensis* BI429 being the most divergent (strain-specific genes, n=33) and the others strain with just a few (BI431, n = 4) or no strain-specific genes (Table 2, Table 3, Supplementary Materials Table S1). The remaining *Thermosipho* spp. genomes have few (BI1063, n = 11) to many unique genes (H17ap60334, n=169) (Supplementary Materials Table S1).

Identification of species-specific genes indicated that each species-level lineage has a common set of genes not detected in the other strain clusters (Table 2). Using 70% IC, the *T. melanesiensis* strains have 424 (+/− 1.2) specific genes (23.0 % of total), while the *T. affectus* and *T. africanus* have 350 (+/− 2)(20.2%) and 650 (+/− 11)(33.4%) cluster specific genes respectively (Table 2). Many of the species-specific genes are hypothetical proteins (26 to 45 %). The *T. africanus* genomes have a larger proportion of species-specific genes in their genome compared to the other two species. Interestingly, when using a less stringent cut-off of 30%, the number of species-specific genes is substantially reduced, e.g. species-specific genes in *T. melanesiensis* BI429 fall to 67 genes (3%), and the proportion of strain specific genes decreases as well. Also here the *T. africanus* genomes have the most species-specific genes, while *T. affectus* isolates have the least (average 5,8 vs 3.0% of total genes per genome). This indicates that most of the species-specific genes have distantly related homologs in the other two species, possibly due to either sequence diversification, or replacement by more distantly related homologs acquired by HGT.

**Table 2.**
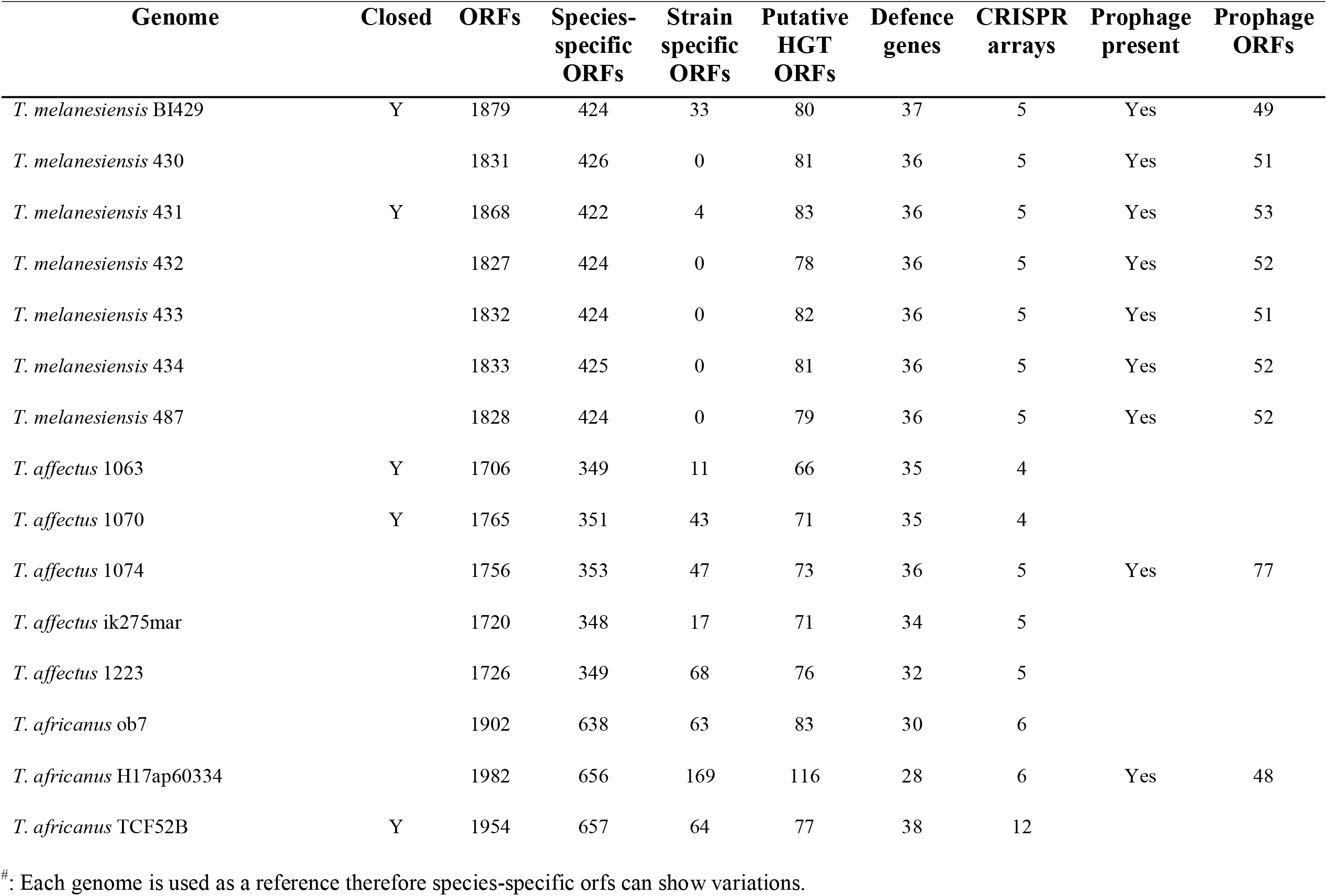
Overview of genome content with a focus on mobile DNA defence systems and mobile elements

### Horizontal gene transfer

Interestingly, in several of the genomes of the *T. africanus* and *T. affectus* isolates we detected blocks of co-localized variable genes (min-max cluster size: 4-31) unique for that genome (Table 3). No such blocks were found between the highly similar *T. melanesiensis* strains. Several of the larger blocks of strain specific genes encode integrated prophages (*T. affectus* BI1074, *T. africanus* H17ap60334) as they could be detected by at least one prophage finding tools (Supplementary information, Supplementary materials Table S1). In addition, when only one of the *T. melanesiensis* genomes was used to find unique genes, we detected one large block consisting of a prophage as well. This prophage was present in all *T. melanesiensis* isolates (Haverkamp et al., manuscript in preparation). The remaining clusters encode genes encoding CRISPR-cas proteins (Table 3) and genes involved in various types of cellular activities. Only a few clusters are dominated by hypothetical proteins (Supplementary materials Table S1).

**Table 3.**
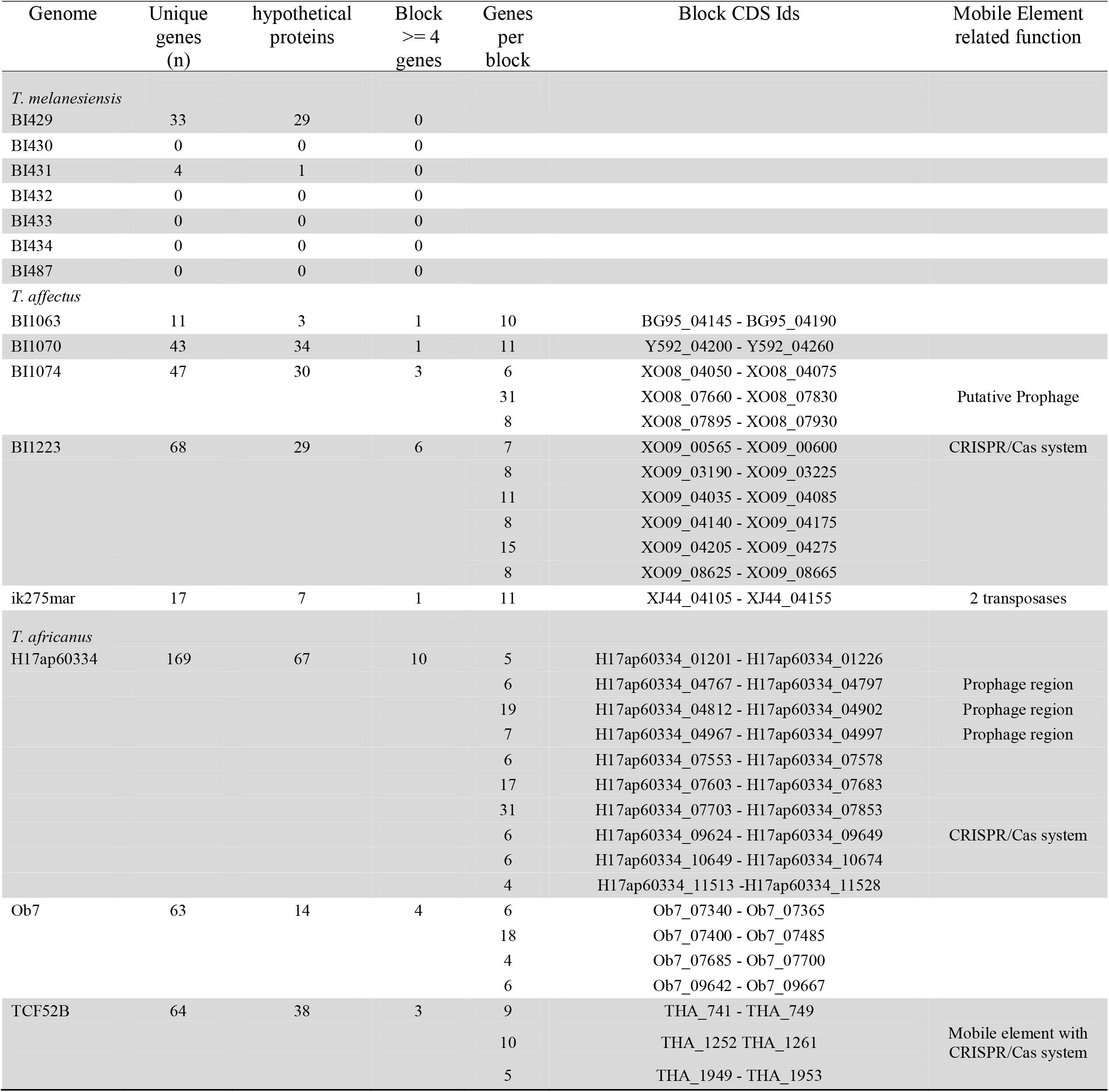
Unique gene counts per genome. Counts were obtained at the IMG database using the phylogenetic profile tool using single genes. The genes of each genome were compared to the genes of all other genomes to identify genes without homologs in the reference genome, with a minimum similarity of 70% and pseudogenes were excluded.

The co-localization of unique genes suggests acquisition by HGT for these clusters. HGTector (Zhu et al. 2014) analysis suggested that between 4.0 % (*T. affectus* BI1063) and 5.8 % (*T. africanus* H17ap60344) of the genes in each genome have been acquired from species not belonging to the Thermotogales by HGT (Table 2, Supplementary materials Table S2). The majority of the putative HGT genes (average ≈85%) showed similarity to genes found in other orders within the Thermotogae, such as the Kosmotogales and the Petrotogales. The remaining genes (average ≈15%) were mainly shared between the *Thermosipho* genomes and genomes of Firmicutes and Euryarchaeota (Supplementary materials Table S2). Within each genome about 50 % of the putative HGT genes belonged to the COG categories: “Carbohydrate transport and metabolism”, “Energy production and conversion”, or were not assigned to a COG category (e.g. Hypothetical genes) (data not shown). The ratio between these three categories was different for the three species, with *T. africanus* having more carbohydrate genes, while *T. affectus* species had more unassigned genes. Interestingly, many of the putative HGT genes could be identified in several *Thermosipho* genomes, which indicates that the common ancestor of the *Thermosipho* isolates acquired most of the putative HGT genes. Only a few putative HGT genes fall into the genome or species-specific gene sets. This is due to the fact that most of these proteins have no high quality blastP match in the genomes in the HGTector database and will therefore not classify as putative HGT.

### Genome-wide comparisons of COG categories

In order to detect functional differences between the three species we compared their genomic content using Clusters of Orthologous Genes (COGs) annotations (Figure 4). This revealed *T. africanus* to have genomes with the highest absolute gene abundances for many COG categories, which is due to the *T. africanus* genomes being larger. However, this effect disappears for most categories when using relative abundances of all COGs (Figure 4A; Supplementary Materials Table S3). The categories H, J, L, R and G had the largest relative abundance differences between the three species (p ≤ 0.01 for H, J and R and p ≤ 0.05 for G and L). Several other categories (B, C, I, O) had highly significant relative abundances differences (p ≤ 0.01), but absolute values were either very low (B), and relative abundance differences were not very large between the species, and within species they were very similar (Supplementary Materials Table S3).

**Figure 4.**
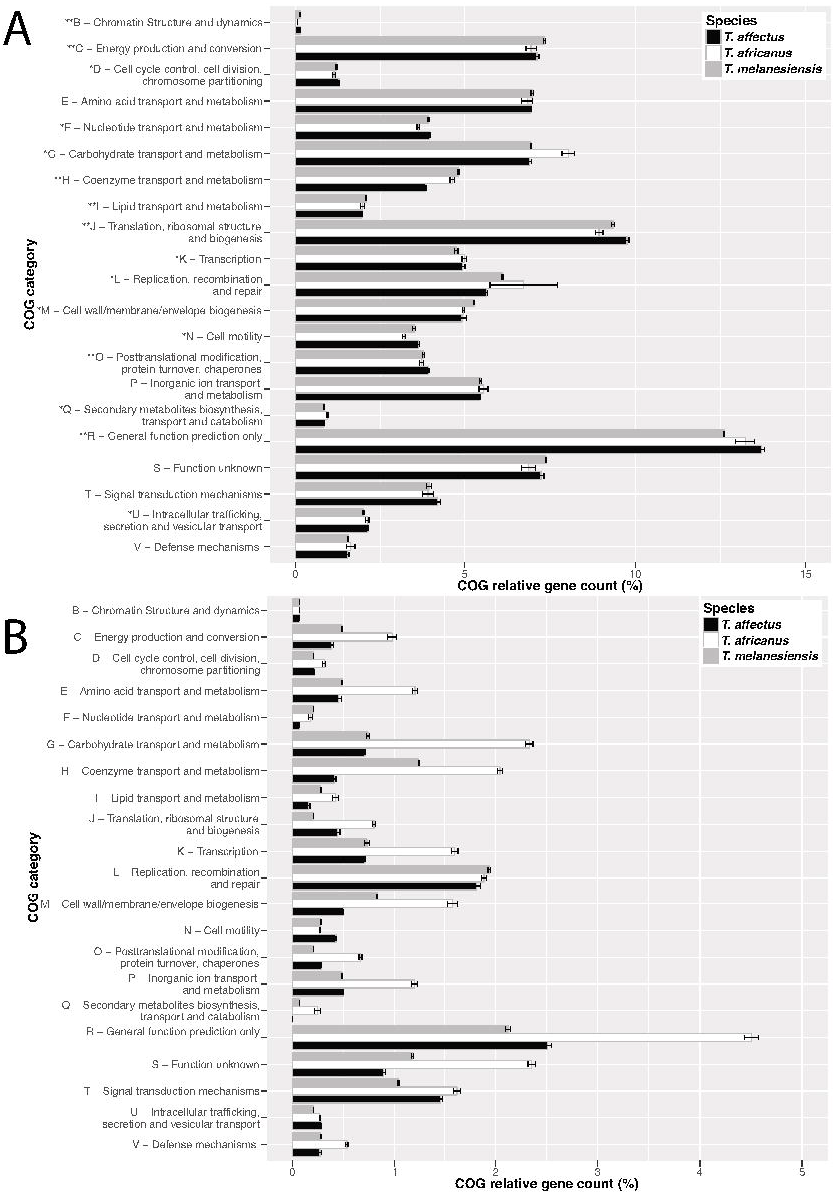
Comparison of average relative COG category gene counts for three *Thermosipho* species. **A)** Complete genome COG category annotations. **B)** Species-specific genes COG annotations. COG gene annotations for each genome were summarized per category and genome counts were standardized using total COG annotation counts per genomes. For the complete genome COG annotations, the Kruskal-wallis test was performed on standardized counts using the species clusters as groups. Categories with significant differences (p-value <=0.01) between the species are indicated with **. The standard error of relative category counts per species, is indicated by the errors bars.

The largest COG category is made up of Category R (General function prediction only), with the *T. affectus* genomes having most genes (Figure 4). Category R is also the largest group among the species-specific genes (Figure 4B). These results are in line with the observation that on average 42.1 %, 49.3 % and 54.8 % (*T. africanus*, *T. affectus* and *T. melanesiensis*) of the species-specific genes lack COG annotations and are hypothetical genes. Interestingly, the *T. africanus* genomes have more genes with COG categories annotations, but they do show more genes with the COG category R. This difference could be caused by the careful manual curation of the TCF52B genome (Nesbø et al. 2009).

The *T. affectus* genomes show a significantly (p ≤ 0.01) lower relative abundance of genes in category H (Coenzyme transport and metabolism category) (Figure 4A). Closer inspection shows that the *T. affectus* genomes are lacking most of the genes (20 out of 22 genes) needed for corrinoid synthesis, except *CobT* and an ATP-binding protein (indicated as ORF). A complete set of corrinoid synthesis genes are found in the genomes of *T. africanus* and *T. melanesiensis*, and are essential for *de novo* vitamine B_12_ synthesis (Swithers et al. 2011) (Supplementary materials Figure S4). Interestingly, the cobalamide salvage pathway gene cluster, which is needed for retrieving incomplete corrinoid molecules from the environment, is present in the *T. affectus* genomes. This gene cluster is, however, missing its *CobT’* gene. This suggests that the orphan *CobT* gene, presumably a remnant from the missing corrinoid cluster, is now functioning in the cobalamide salvage pathway (Supplementary materials Figure S4).

Large differences among the genomes were also seen for COG category G (Carbohydrate transport and metabolism). In agreement with this, phenotypic differences in carbohydrate metabolism is one of the main other features, to distinguish between the three species (Podosokorskaya et al. 2011). Also for this category, we find that the *T. africanus* genomes have relatively more genes than the other two species (Figure 4A). This difference is even more pronounced for species-specific genes (Figure 4B), where *T. africanus* genomes have more genes present in this category (Supplementary Materials Table S3). A screening of the genomes using PFAM annotations and the carbohydrate database dbCAN (Yin et al. 2012), showed a similar pattern as with the COG annotations (Supplementary information; Supplementary materials Table S4). The *T. affectus, T. melanesiensis and T. africanus* genomes contain on average: 16-17, 20 and 21-26 genes respectively that are involved in breakdown of carbohydrates (Supplementary materials Table S4). Moreover, the families, containing enzymes involved in breakdown of various beta-linked oligo-and polysaccharides (eg. cellulose, xylan, laminarin, lichenan, mannans and chitin) were found exclusively among the representatives of *T. africanus*. This shows, in line with the COG analysis, that the *T. africanus* species might be more versatile with regard to carbohydrate uptake and metabolism.

**Table 4.**
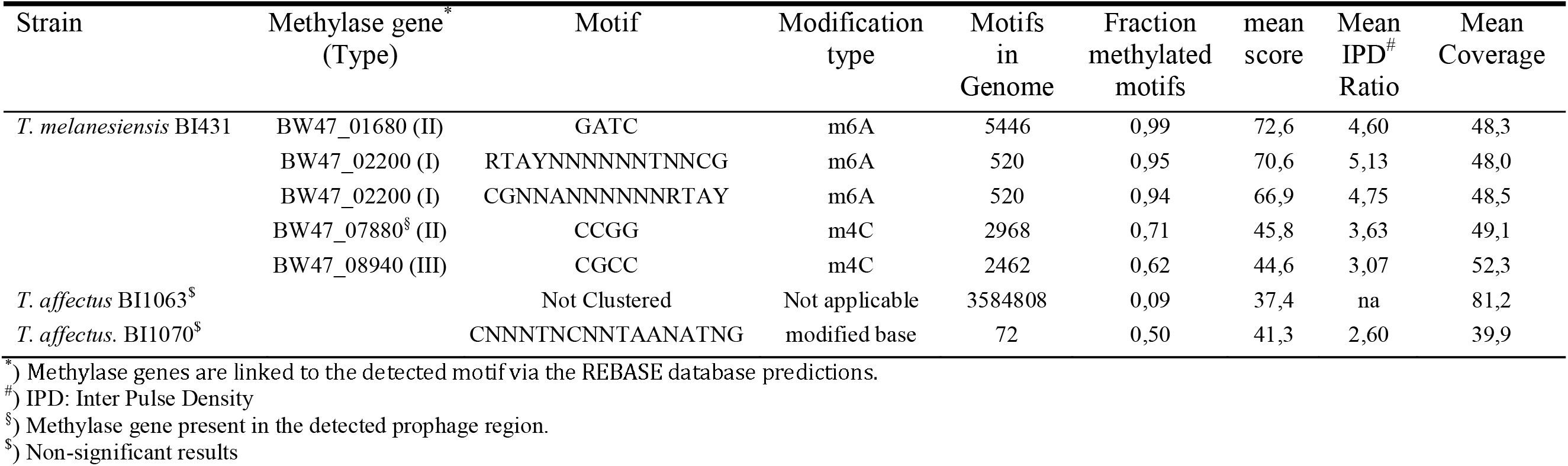
Overview of modification and motif analysis for three *Thermosipho* strains using Pacbio sequencing. Modification and motif analysis was performed using the RS_modifcation_and_motif_analysis.1 pipeline using the SMRT sequencing subreads and as a reference the non-closed quiver polished HGAP assembly for each genome. Bases with m6A methylation were identified along the entire chromosome sequences of *Thermosipho melanesiensis* 431.

When we compared the COG categories for the species-specific genes we found even larger differences for many categories (Figure 4B). Since the *T. africanus* genomes have almost twice as many species-specific genes compared to the other two species, they also have proportionally more species-specific genes in most categories. For instance, for COG category L (Replication, recombination and repair) we find large variation in the number of genes among the three *T. africanus* strains, with TCF52B having relatively more of these genes compared to the other genomes (8.7 % vs 5.7%). Examination of the genes assigned to this category revealed that this difference is mainly due to the presence of 18 copies of transposases in the TCF52B genome.

### Defense genes

Above we identified several interesting features of the *Thermosipho* genomes, where some of the genomes have a large set of unique genes, contain large mobile elements, or have many putative HGT genes. This suggests that the defense mechanisms against mobile DNA might differ among the isolates. We therefore screened the annotated genomes for the presence of COGs that are related to defense mechanisms (Makarova et al. 2011), resulting in a list of 44 COGs that could be classified into three clusters, restriction-modification (RM) systems genes (12), CRISPR-cas genes (18) and other COG annotations (14) (Figure 5).

**Figure 5.**
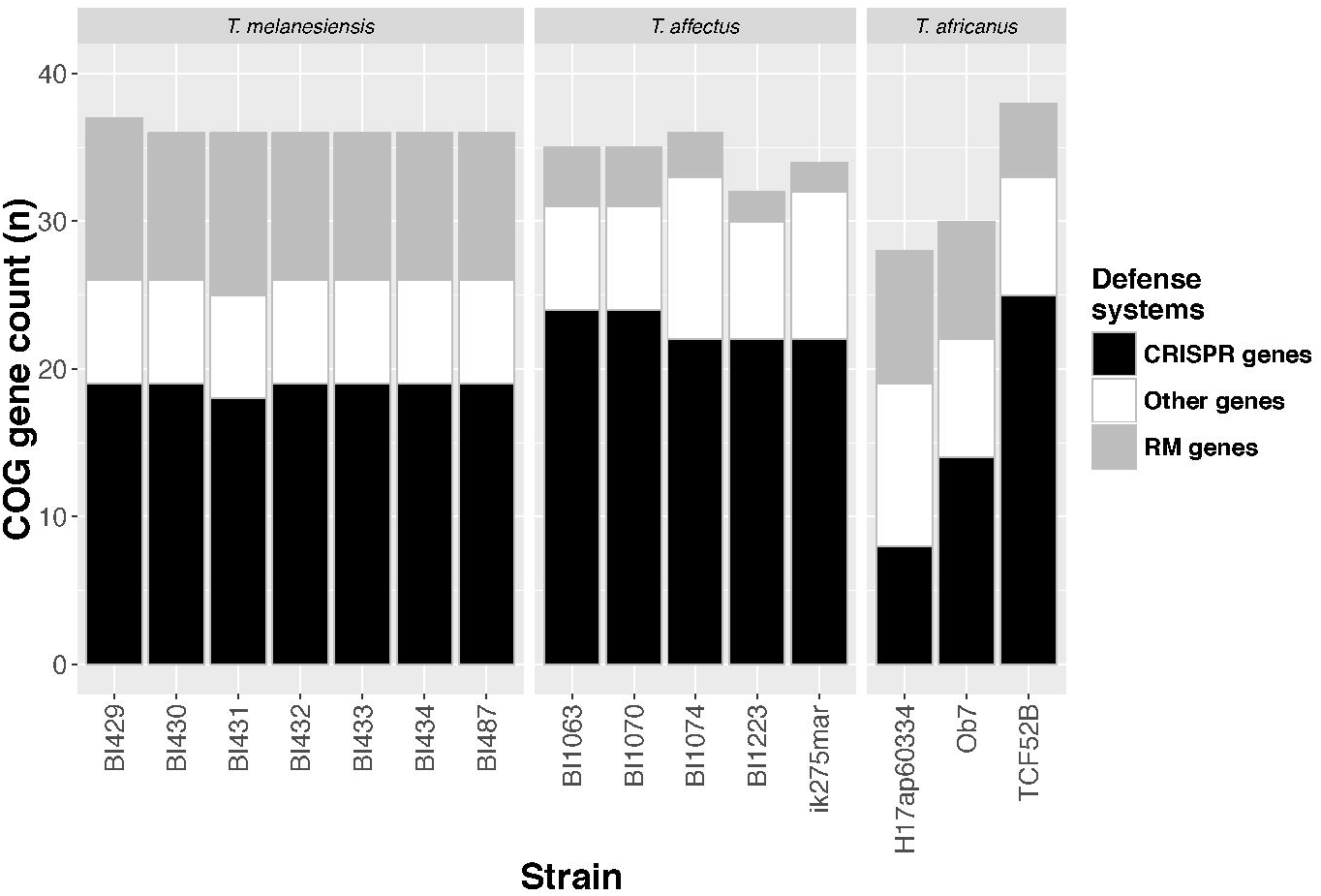
Mobile DNA defense related COG annotation counts for 15 *Thermosipho* isolates. A total of 38 COGs were identified using a list of COGs related to defense genes (Makarova et al. 2011). The counts of the identified COGs were summarized into three groups: Restriction-modification systems COGs, CRISPR-cas associated COGs, and other COGs. The strains are grouped per *Thermosipho* species.

Overall, the *T. affectus* genomes and *T. africanus* TCF52B have the most CRISPR-cas genes (Figure 5). Specifically, we observed that all the *Thermosipho* genomes contain *cas10*/*crm2* (COG1353), which indicates the usage of the type III CRISPR-cas system (Supplementary materials Table S5) (Makarova et al. 2015). The *cas10*/*crm2 (*COG1353) annotated genes are present at multiple loci in *T. affectus* and *T. melanesiensis* strains, but only one copy is found in *T. africanus* isolates. For the proper functioning of the type III system, *crm6* (COG1604) is needed. However, this gene is missing in *T. africanus* H17ap60334, suggesting an incomplete and possibly non-functioning CRISPR-cas system (Makarova et al. 2015). Furthermore, the *T. africanus* H17ap60334 genome has a lower number of CRISPR-cas genes compared to the other isolates (Figure 5; Supplementary materials Table S5). All the *T. melanesiensis* strains (except BI431) and *T. africanus* strain TCF52B also carry the type I CRISPR-cas system (marker gene: *cas3* (COG1203)). This gene is not detected in the *T. affectus* or *T. africanus* Ob7 and H17ap60334 genomes, suggesting there are differences in the mechanism of the CRISPR-cas system both within and between the *Thermosipho* isolates / species.

We also detect differences in RM systems-gene content. The *T. affectus* genomes contain few RM system genes and the *T. melanesiensis* have the most (Figure 5; Supplementary materials Table S5). For a functioning RM system both the methylase and restriction enzymes are needed. The restriction enzymes are absent from the *T. affectus* genomes, which suggest that this species does not possess a functional RM-system (Supplementary materials Table S4). In contrast, we detected both type I, II and III restriction enzymes and DNA methylases in the *T. melanesiensis* genomes. In the *T. africanus* isolates a complete type II system is present in the genomes of strains Ob7 and H17ap60334. In addition, all the *T. africanus* strains have a complete type III RM system (Supplementary materials Table S5).

In agreement with the distribution of RM-systems, Pacbio sequencing of *T. melanesiensis* BI431 and *T. affectus* BI1063 and BI1070 revealed that strain BI431 was methylated, while the two *T. affectus* genomes lacked methylation. The *T. melanesiensis* BI431 methylation sites detected matched those predicted in the Rebase restriction database (Roberts et al. 2015); both methyl-6-adenosine (m6A) and methyl-4-cytosine (m4C) methylation were detected along the entire BI431 genome. The dominant methylation sites indicate that methylation was mainly due to the type II and III systems (Table 4). We did not perform Pacbio sequencing of the *T. africanus* genomes, but the presence of a type-III methylase enzyme suitable for a functional RM-system suggests the possibility of a methylated genome.

### CRISPR array variation

Between four and six CRISPR arrays were detected in all the genomes, except for *T. africanus* TCF52B, which contains 12 arrays (Table 2, Table 5, supplementary materials table S6). The total number of spacers detected in all genomes was 1709, with spacer count varying between 54 (*T. affectus* BI1223) to 321(*T. africanus* TCF52B). All the spacers could be clustered into 1001 different clusters, indicating that there are both shared and unique spacers. For instance, the highly similar *T. melanesiensis* genomes all have 5 CRISPR arrays, with a total of 681 spacers that form 92 clusters that are present in all *T. melanesiensis* genomes (Supplementary material Figure S5A). The only difference observed between these genomes was strain BI429 missing four spacers found in the other genomes. Interestingly, six of the 92 clusters have spacers from different arrays (Supplementary material Figure S5A).

**Table 5.**
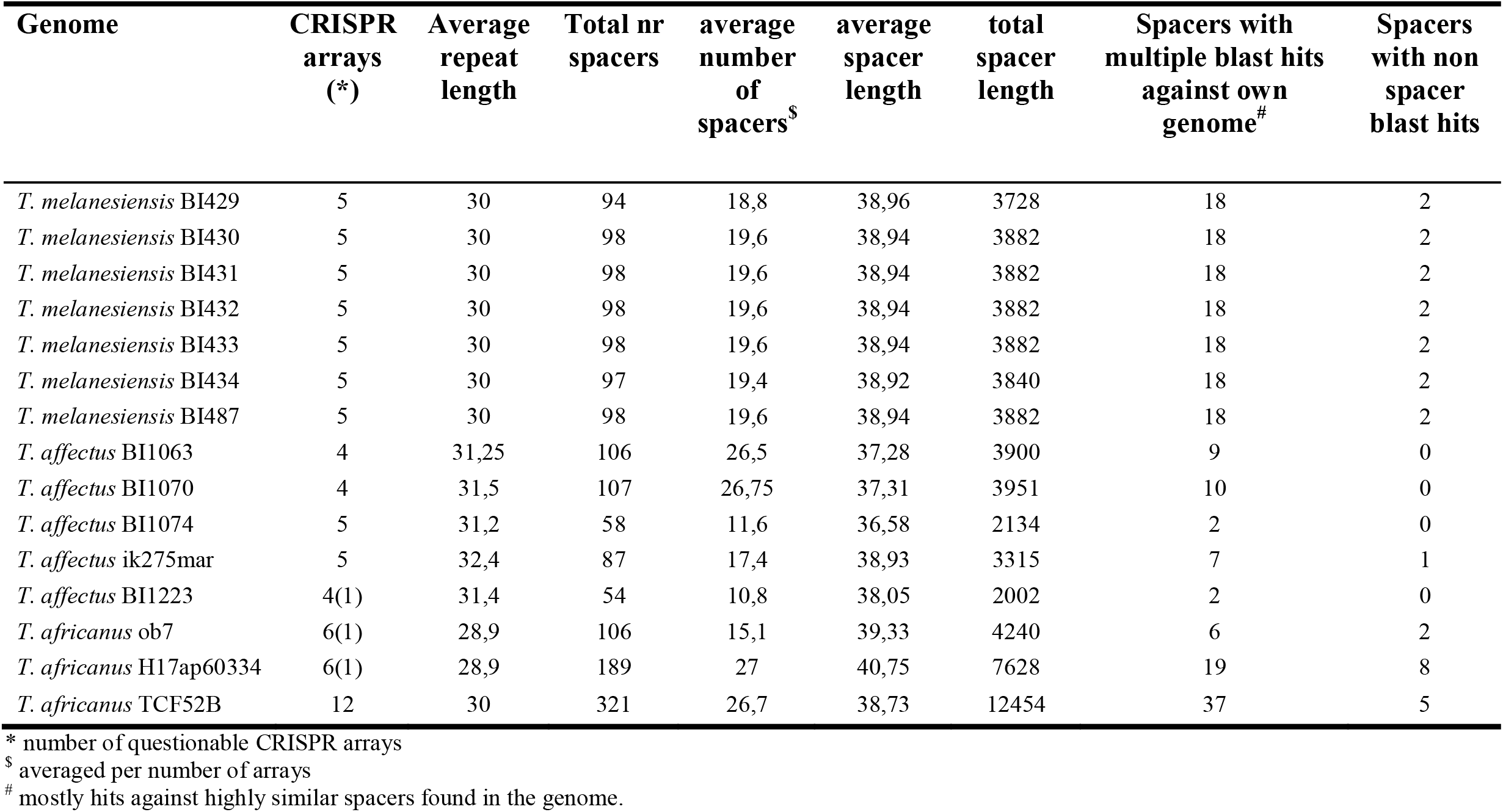
CRISPR array comparison of 15 *Thermosipho* genomes. CRISPR arrays were detected using CRISPRfinder (Grissa et al. 2007). Spacers were extracted and blasted against the own genome to determine if there were other similar sequences present in the genome. Most spacers that showed hits, matched other spacers found in one of the CRISPRarrays of the genome.

| Genome | CRISPR<br>arrays<br>(*) | Average<br>repeat<br>length | Total nr<br>spacers | average<br>number<br>of<br>spacers<sup>\$</sup> | average<br>spacer<br>length | total<br>spacer<br>length | Spacers with<br>multiple blast hits<br>against own<br>genome<sup>#</sup> | Spacers<br>with non<br>spacer<br>blast hits |
| --- | --- | --- | --- | --- | --- | --- | --- | --- |
| *T. melanesiensis* BI429 | 5 | 30 | 94 | 18,8 | 38,96 | 3728 | 18 | 2 |
| *T. melanesiensis* BI430 | 5 | 30 | 98 | 19,6 | 38,94 | 3882 | 18 | 2 |
| *T. melanesiensis* BI431 | 5 | 30 | 98 | 19,6 | 38,94 | 3882 | 18 | 2 |
| *T. melanesiensis* BI432 | 5 | 30 | 98 | 19,6 | 38,94 | 3882 | 18 | 2 |
| *T. melanesiensis* BI433 | 5 | 30 | 98 | 19,6 | 38,94 | 3882 | 18 | 2 |
| *T. melanesiensis* BI434 | 5 | 30 | 97 | 19,4 | 38,92 | 3840 | 18 | 2 |
| *T. melanesiensis* BI487 | 5 | 30 | 98 | 19,6 | 38,94 | 3882 | 18 | 2 |
| *T. affectus* BI1063 | 4 | 31,25 | 106 | 26,5 | 37,28 | 3900 | 9 | 0 |
| *T. affectus* BI1070 | 4 | 31,5 | 107 | 26,75 | 37,31 | 3951 | 10 | 0 |
| *T. affectus* BI1074 | 5 | 31,2 | 58 | 11,6 | 36,58 | 2134 | 2 | 0 |
| *T. affectus* ik275mar | 5 | 32,4 | 87 | 17,4 | 38,93 | 3315 | 7 | 1 |
| *T. affectus* BI1223 | 4(1) | 31,4 | 54 | 10,8 | 38,05 | 2002 | 2 | 0 |
| *T. africanus* ob7 | 6(1) | 28,9 | 106 | 15,1 | 39,33 | 4240 | 6 | 2 |
| *T. africanus* H17ap60334 | 6(1) | 28,9 | 189 | 27 | 40,75 | 7628 | 19 | 8 |
| *T. africanus* TCF52B | 12 | 30 | 321 | 26,7 | 38,73 | 12454 | 37 | 5 |

\* number of questionable CRISPR arrays
<sup>\$</sup> averaged per number of arrays
<sup>#</sup> mostly hits against highly similar spacers found in the genome.

The other two species show much more diversity in comparison to *T. melanesiensis*, within the spacer sequence content between the strains of the different species (Supplementary material Figure S5 B and C). The *T. affectus* spp. CRISPR array spacers (n=412) fall into 366 clusters, while the *T. africanus* spp. CRISPR array spacers (n= 616) form 543 clusters.

Comparing spacer sequences to the host genome (excluding CRISPR arrays, blastN e-value cut-off: 1.0^-5^) revealed a small fraction of the spacers with matches within its host genome (percentage identity 88-100%) (Table 5). The *T. affectus* genome spacers did not match with any region in the host genome. For the *T. melanesiensis* genomes one spacer (array 2, spacer 5) matched one gene (e.g. *T. melanesiensis* BI429: Tmel_1466: hypothetical protein) in all genomes (81.8% identity, 6 SNPs). This gene is located within a prophage element consisting of 50 protein coding genes (Tmel_1439: Tmel_1486) (Haverkamp et al., in prep). The relatively low similarity between the spacer and the gene sequence could suggests that this gene is not or no longer a target sequence, but we can not exclude it (Cady & O'Toole 2011).

For the *T. africanus* genomes we detected one CRISPR spacer, shared by all three genomes, which has a non-perfect match to the same genomic region. The spacer matches a phospholipase / carboxylesterase gene (THA_1282 in strain TCF52B) with 89-92% identity. For *T. africanus* H17ap60334 we find three additional spacers with perfect matches in two of its gene; two spacers target H17ap60334_04822 and one targets H17ap60334_04912. Closer examination revealed that also these genes are part of a predicted prophage region (Haverkamp et al., in prep.; Supplementary information).

Finally, we searched NCBI’s non-redundant nucleotide database for matches to the *Thermosipho* spacer sequences. This identified two identical 44 bp spacers, one in *T. africanus* H17ap60334 (array 2, spacer 7) and one in *T. africanus* TCF52B (array 5, spacer 20) that matched a sequence in the genome of *Pseudothermotoga elfii* DSM 9442 (genbank refseq ID: NC_022792). The identified region in that genome (bp 200095 – 200125) is a CRISPR spacer sequence of 38bp, which is identical to the *Thermosipho* spacer for 31 bases. Interestingly, the first six bases of the *Pseudothermotoga* spacer has no match to the *Thermosipho* H17ap60334 and TCF52B spacer, while the last 12 bases of the *Thermosipho* spacers have no match to the *Pseudothermotoga* spacer. This suggests the spacers in both species were acquired independently, but that they match a similar sequence.

## Discussion

Bacterial genome stability and evolution are under influence of HGT, which, among other things, may influence how quickly a new species will arise (Darmon & Leach 2014). For instance, low levels of DNA exchange, within species, will allow mutations to arise quicker than homologous recombination can repair it leading to speciation events, while high levels of within species DNA exchange, prevents strains from accumulating enough mutations to differentiate (Lawrence & Retchless 2009). High levels of HGT have previously been observed for Thermotogae bacteria, in particular the *Thermotoga* genus, which is found in high temperature ecosystems, comparable to those where *Thermosipho* occurs (Nelson et al. 1999; Zhaxybayeva et al. 2009; Nesbø et al. 2015). Some of these habitats can be very different, e.g. marine hydrothermal vents and deep subsurface oil reservoirs. It is however unclear, how habitat-range influences genome structure and content, levels of homologous recombination and HGT (and *vice versa*).

Here we shed light on these questions by analyzing multiple isolates of the Thermotogae genus *Thermosipho*. The isolates were obtained from various ecosystems across the globe (Table 1) and fall into 3 different lineages or species: *T. melanesiensis*, *T.africanus* and *T. affectus. T. melanesiensis* and *T. affectus* isolates were only obtained from hydrothermal vents from one specific geographical region, while *T. africanus* members have been obtained from a wide range of geographical and ecologically different environments (oil reservoirs, hydrothermal vents and terrestrial hot springs). This implies that the genus *Thermosipho* has specialist and generalist species, with the latter being present in a wider range of ecosystems.

Although the above observation may be partly due to sampling and cultivation bias, comparison with 16S rRNA genes from environmental samples (Dahle et al. 2008; Nakagawa:2006jr; Stevenson et al. 2011; Smith et al. 2017) also supports this view, where the two lineages restricted to hydrothermal vents from a distinct branch in the 16S rRNA tree (Figure 1). The fact that these lineages are restricted to hydrothermal vents and appear to have more limited geographic ranges, suggests that these are more specialized species in comparison to species found on the other 16S rRNA tree branch. Interestingly, in addition to oil reservoirs, *Thermosipho* sp. have also been detected in subsurface crustal fluids Smith et al., (2016) and their amplicon 16S rRNA sequences align best with species of the “generalist” branch of the 16S rRNA tree (Data not shown). These results suggest that *Thermosipho* lineages are widespread in the subsurface and not restricted to oil reservoirs or hydrothermal vents.

Another representative of the *Thermotogae* species, where ecology is suggested to be important for differentiation, is *Thermotoga maritima*. For this lineage, similar ecology was found to be more important than close geographical distance to maintain highly similar genomes through extensive gene flow (Nesbø et al. 2015); genomes from the same type of environment (i.e. oil reservoir or marine vent) were more similar to each other than geographically close genomes from different types of environments (Nesbø et al. 2015).

As in *T. maritima* we found that within *Thermosipho* sp. gene flow appears to be important for the maintenance of the three distinct lineages or species, as we observe high levels of recombination within the three lineages investigated. However, we also see a clear differentiation of the three species with no recombination between lineages. These results were further supported by the pangenome analysis of the genus *Thermosipho* (Figure 3B). Each lineage, or species, contained a set of species-specific genes that were at best only distantly related to genes in the other lineages. These genes mark the species boundaries and possibly contribute to niche differentiation between the different *Thermosipho* species. This was especially clear from the analysis of genes involved in carbohydrate and coenzyme metabolism (COG category G and H).

These carbohydrate and coenzyme metabolism genes are important for energy production and central metabolism and could play a role in the niche differentiation of the *Thermosipho* species. For instance, the *T. affectus* genome contains several genes with a dependency for vitamin B12 (e.g. B12 dependent methionine synthase, ribonucleotide reductase Class II)(Swithers et al. 2011). However, the absence of most genes needed for the corrinoid synthesis (needed for de-novo Vitamin B12 synthesis) (Coenyzme category) in *T. affectus* suggests that this organism can only rely on the recovery of Vitamin B12 precursor molecules (Supplementary materials Figure S4). It is unclear how this affects its physiology and ecology. It will however, certainly affect the type of environments it can inhabit and its role and interaction with other species in the communities they are part of. Likewise, *T. africanus* has more genes involved in carbohydrate metabolism, which suggests that this species is more diverse with respect to which carbohydrate molecules it can utilize. This seems to be in line with the literature, (Podosokorskaya et al. 2014; Dipasquale et al. 2014) and it could be partly responsible for its wider distribution in different ecosystems.

It is notable, that the species with the largest, and also most variable genomes, have the widest distribution, both geographically and ecologically. However, the difference in genomic variation *within* three lineages is probably also partly due of the fact that the *T. africanus* isolates were sampled from three different geographically separated populations, while the *T. melanesiensis* isolates all originated from a single population sampled at the same time from the same ecosystem. The *T. affectus* genomes originated from two geographically close sites and show intermediate levels of variation (Table 1). These results suggest that isolation by distance is an important factor for differentiation in *Thermosipho* sp., as found in other thermophiles {Mino:2017gj}. However, since *T. affectus* and *T. melanesiensis* have not been detected at other locations than the isolation sites described here, this may be an entirely academic question for these species.

### Dealing with invasive DNA

Phages and transposons are known vectors of “foreign” DNA with possible deleterious effects, therefore limiting their integration and activity is an essential survival strategy for many, if not all, bacterial species (Stern & Sorek 2011). Several mobile elements were detected in the *Thermosipho* genomes. For instance, *T. africanus* TCF52B carries 18 copies of a transposase (COG2801). Moreover, one *T. africanus* (H17ap60334), one *T. affectus* (BI1074) and all the *T. melanesiensis* strains have a prophage integrated into their genomes (Haverkamp et al., unpublished). The presence of prophages in Thermotogales species has earlier only been described in the thermophile *Marinitoga piezophila*, and analysis of the phage (named MPV1) genome suggested it has played an important role as a HGT vector (Lossouarn et al. 2015). A large fraction of its genes show a close phylogenetic relationship with Firmicutes, the Thermotogae’s main HGT partner (Zhaxybayeva et al. 2009). The identification of multiple phages in *Thermosipho* genomes further supports the idea that phage particles could be the general intermediates in gene flow within and between species in the subsurface (Labonté et al. 2015).

Nonetheless, phages can be detrimental for prokaryotic cells such as *Thermosipho*, so mechanisms are in place in this genus to intercept incoming foreign DNA. These mechanisms consist of the RM and the CRISPR-cas system, that both play a role in intercepting invasive DNA. The presence of CRISPR-cas genes, which are involved in prokaryotic immunity, among the species-and strain specific genes is no surprise since they have been identified before on mobile elements such as transposons and phages (Sebaihia et al. 2006; Seed et al. 2013). One example of this is a mobile element in *T. africanus* TCF52B flanked by transposase recognition sites, which contains CRISPR-cas genes (Nesbø et al. 2009). This further indicates that mobile elements are not always deleterious for the receiver, but they can be a source of novel beneficial genetic material.

The comparative genome analysis shows that there are clear differences between the three *Thermosipho* species in the completeness of the systems, which may have implications for acquiring novel DNA via HGT or recombination. For instance, all isolates have the CRISPR-cas system, but the number of genes per genome related to this system differs, and in *T. africanus* H17ap60334 the CRISPR-cas system may not be complete enough to be functional. On the other hand, in contrast to the two other species the RM system is incomplete in *T. affectus* species and likely not functional, indicating that *T. affectus* isolates only rely on the CRISPR-cas system for defense against foreign DNA.

The lack of a working RM system was confirmed through PacBio sequencing of strains *T. affectus* BI1063 and *T. affectus* BI1070 that showed no methylation. Pacbio sequencing of *T. melanesiensis* BI431 revealed a methylated genome supporting the presence of a working RM system. The genome of the isolate and the integrated prophage were methylated. One of the RM systems detected in *T. melanesiensis* BI431 is located in the integrated prophage, but it is not the only system that is active. Both host and prophage methylases are expressed and methylation is detected at sites specific for those methylases (Table 4). The presence and activity of a methylase in the prophage genome is a well know evasive mechanism deployed by phages to escape the host RM system (Stern & Sorek 2011).

That the host RM system is active suggests that *T. melanesiensis* strains repeatedly encounter mobile DNA / phages in their natural environment. This is also suggested by the low similarity of one of the CRISPR array spacers with the prophage genome, which implies that considerable divergence has occurred since the creation of the spacer and the integration of the detected prophage. If the sequence match had been perfect, it would have been likely that the prophage could not have integrated since the CRISPR machinery would have stopped it at the door. Another interesting observation was the presence of a spacer, in both *T. africanus* strains TCF52B and H17ap60334, similar but not identical to a spacer in the genome of *Pseudothermotoga elfii* DSM 9442. This suggests the presence in the subsurface of a broad range phage infecting both species.

Taken together, the observations of the diversity of CRISPR-cas systems, RM systems, and mobile elements (i.e. prophages and transposons) in *Thermosipho*, indicate that mobile DNA, such as phages and transposons, frequently interact with these bacterial genomes in the subsurface. Recently, it was shown that phages are dominant players in the subsurface realm, influencing ecosystem function by interacting with their prokaryotic hosts (Engelhardt et al. 2015). It also implies that mobile DNA plays an important role in shaping *Thermosipho* populations and species in the subsurface.

### Conclusions

Here we compared the genomes of fifteen different *Thermosipho* isolates and found that they belong to three clearly separated lineages or species: *T. affectus, T. africanus* and *T. melanesiensis*. Recombination detection showed very little interspecies gene flow, but high intraspecies gene flow. This finding was supported by the presence of large groups of species-specific gene sets, reflecting the metabolic differences between the species. The species-specific genes sets may further be responsible for differences in distribution of these species in various ecosystems such as hydrothermal vents and petroleum reservoirs. Our observations suggest that genome similarity in this genus decreases with increasing distance. However, additional genome sequences of *Thermosipho* isolates from different locations and lineages within this genus are needed in order to confirm this. Our analysis also revealed that the presence of prophage elements is not uncommon in these thermophiles. It also showed that each of the species has a different capacity for dealing with incoming DNA from phages and other mobile elements, which also affects intra-species gene flow. The presence of multiple prophage elements in several *Thermosipho* isolates suggests that the high fraction of horizontally acquired genes is possibly due to ongoing warfare between bacteria and phages in the subsurface. Finally, the presence of similar CRISPR spacers, in multiple species and isolates indicates that phages in the subsurface have broad host ranges allowing for inter-species gene flow. That could ultimately explain why many genes in Thermotogales genomes show high similarity with genes found in Archaea or Firmicutes genomes.

## Acknowledgements

The work of THAH and CLN, was supported was funded by the Norwegian Research Council (award 180444/V40). JL and CG were supported by Agence Nationale de la Recherche (ANR) (ANR-12-BSV3-OO23-01). OAP and IVK were supported by the Russian Science Foundation (RSF) grant # 16-14-00121. We thank the Norwegian Sequencing Centre for sequencing and support with the bioinformatics analysis. We thank Ifremer and chief scientists of the French oceanographic cruises “Biolau” (1989), “Marvel” (1997) and “Atos” (2001). Strains are available under request at the ‘Souchothèque de Bretagne’ (catalogue LMBE) culture collection (http://www.univ.brest.fr/souchoteque/collection+LM2E).

